# Tuning the resorption-formation balance in an *in vitro* 3D osteoblast-osteoclast co-culture model of bone

**DOI:** 10.1101/2022.08.04.502780

**Authors:** Stefan J.A. Remmers, Freek C. van der Heijden, Bregje W. M. de Wildt, Keita Ito, Sandra Hofmann

## Abstract

The aim of the present study was to further improve an *in vitro* 3D osteoblast (OB) – osteoclast (OC) co-culture model of bone by tuning it towards states of formation, resorption, and equilibrium for their future applications in fundamental research, drug development and personalized medicine. This was achieved by varying culture medium composition and monocyte seeding density, the two external parameters that affect cell behavior the most. Monocytes were seeded at two seeding densities onto 3D silk-fibroin constructs pre-mineralized by MSC-derived OBs and were co-cultured in one of three different media (OC stimulating, Neutral and OB stimulating medium) for three weeks. Histology showed mineralized matrix after co-culture and OC markers in the OC medium group. Scanning Electron Microscopy showed large OC-like cells in the OC medium group. Micro-computed tomography showed increased formation in the OB medium group, equilibrium in the Neutral medium group and resorption in the OC medium group. Culture supernatant samples showed high early TRAP release in the OC medium group, a later and lower release in the Neutral medium group, and almost no release in the OB medium group. Increased monocyte seeding density showed a less-than-proportional increase in TRAP release and resorption in OC medium, while it proportionally increased TRAP release in Neutral medium without affecting net resorption. The 3D OB-OC co-culture model was effectively used to show an excess of mineral deposition using OB medium, resorption using OC medium, or an equilibrium using Neutral medium. All three media applied to the model may have their own distinct applications in fundamental research, drug development, and personalized medicine.

## Introduction

Bone growth and homeostasis is regulated by bone forming osteoblasts (OBs), bone resorbing osteoclasts (OCs), and regulating osteocytes. These cells tightly regulate bone mass, bone strength and bone structure to continuously meet the requirements placed upon bone tissue. Disturbances to this balance can lead to diseases such as for example osteoporosis. While many treatment options are available for osteoporosis that can delay the progression of the disease, there is currently no cure to this degenerative disease [1,2]. Many of the biochemical actors in bone remodeling have been identified [2–4], but much remains to be learned on the precise nature of the biochemical interplay orchestrating bone remodeling before osteoporosis can be treated or even cured as opposed to merely slowing down the degenerative nature of the disease.

Accurate, scalable, and translatable experimental models are needed to further study the mechanisms underlying bone remodeling. Options such as animal models are expensive, time consuming and far from scalable. They raise ethical concerns and often lead to poor translation from pre-clinical trials to clinical use [5–7]. In contrast, *in vitro* cell-culture models do not share those ethical concerns, can be developed into high-volume tests, and can use cells of various origins, including healthy human donors or even cells from patients suffering from bone diseases [8,9].

*In vivo* bone remodeling is a three-dimensional process where the cells from the basic multicellular unit [10] deposit and resorb three-dimensional volumes of bone tissue. This makes the use of 2D models [11,12] less appealing for studying bone remodeling, especially when cells in 2D monolayer often respond differently than in a 3D environment [13,14]. Although usually easier to obtain, animal cells can respond differently than human cells [9], possibly introducing errors due to interspecies differences. Consequently, to study remodeling and to quantify effects on both resorption and formation within the same model system, ideally a 3D environment is used in which at least both human OBs and OCs are co-cultured simultaneously [15] and can interact freely with each other through both cell-cell contact and paracrine signaling [2].

3D OB-OC co-culture models exist [16,17] where resorption and formation are studied using destructive techniques such as using for example Alizarin Red mineralized nodule staining [18] or Scanning [19] and Transmission [20] electron microscopy for resorbed surface metrology. Longitudinal monitoring of bone remodeling offers the advantages of measuring changes within the same constructs over time and localizing where formation and resorption events take place within constructs. Longitudinal monitoring using micro-computed tomography (μCT) has been shown in animal models before [21–23], and was recently applied to monitor scaffold mineralization by mesenchymal stromal cell (MSC)-derived OBs [24] and subsequent OC resorption [25]. In these studies, the crosstalk occurring in the cultures was biochemically ‘overruled’ to obtain maximal formation and resorption. In a healthy *in vivo* situation, crosstalk between cells results in an equilibrium between formation and resorption, while bone diseases manifest as a disbalance between formation and resorption.

To effectively apply this model to applications such as personalized medicine, drug testing and fundamental research, the model should allow the measurement and visualization of the effects of an external stimulus (e.g. a drug or biochemical compound) on cell activity, formation and resorption. This requires further improvement of the cellular response of the 3D co-culture model on these outcome measures. The two characteristics most suitable for tuning the OC activity in this model are 1) co-culture medium and 2) monocyte seeding density. That is because these characteristics can easily be manipulated and are expected to have a large impact on OC differentiation and function. Culture medium and its components determine to a large extent the proliferation, differentiation and gene expression behavior of cells [26,27], while seeding density determines the extent of intercellular communication, maturation, and in the case of monocytes in particular, also the fusion towards OCs [28].

The aim of this study was to improve the earlier developed co-culture model by investigating how to steer the response of the co-culture model towards and away from resorption, formation, and equilibrium.

## Materials and Methods

### Materials

This study was reviewed and approved by the ethics review board of the European Research Council (ERC) before the study began. Human bone marrow was commercially purchased from Lonza (Walkersville, MD, USA), collected under their institutional guidelines and with written informed consent. A human buffy coat was obtained from Sanquin (Eindhoven, Netherlands) after review and approval of the study by the Sanquin ethics review board. The buffy coat was collected by Sanquin under their institutional guidelines and with written informed consent per Declaration of Helsinki. Antigen retrieval citrate buffer, RPMI-1640 medium, poly-L-lysine coated microscope slides and SnakeSkin Dialysis tubing (3.5 kDa molecular weight cut-off) were from Thermo Fisher Scientific (Breda, The Netherlands). Disposable biopsy punches were from Amstel Medical (Amstelveen, the Netherlands). Trypsin-EDTA (0.25 %) was from Lonza (Breda, The Netherlands). Dulbecco’s modified Eagle media (DMEM low glucose + high glucose), non-essential amino acids (NEAA) and antibiotic/antimycotic (anti-anti) were from Life Technologies (Bleiswijk, The Netherlands). Fetal Bovine Serum (FBS, batch F7524-500ML / lot BCBV7611) was from Sigma Aldrich / Merck. Lymphoprep™ was from Axis-Shield (Oslo, Norway). MACS^®^ Pan Monocyte Isolation Kit was from Miltenyi Biotec (Leiden, the Netherlands). Recombinant human basic fibroblast growth factor (bFGF), macrophage colony stimulating factor (M-CSF) and receptor activator of nuclear factor kappa-B ligand (RANKL) were from PeproTech (London, United Kingdom). *Bombyx mori L*. Silkworm cocoons were from Tajima Shoji Co., LTD. (Yokohama, Japan). Antibody Integrin β3 (Orb248939, Mouse, 1:100) was from Biorbyt (Cambridge, United Kingdom). Antibody TRAP (Sc-30833, Goat, 1:100) was from Santa-Cruz Biotechnology, Inc. (Heidelberg, Germany). Antibody Alexa488 (715-545-150, Donkey-anti-Mouse IgG (H+L), 1:300) was from Jackson ImmunoResearch (Cambridgeshire, United Kingdom). Antibody Alexa488 (A11055, Donkey-anti-Goat IgG (H+L), 1:300) was from Molecular Probes (Eugene, OR, USA). Thin bleach was from the local grocery store. All other substances were of analytical or pharmaceutical grade and obtained from Sigma Aldrich / Merck (Zwijndrecht, The Netherlands).

### Methods

#### Monocyte isolation

A human peripheral blood buffy coat from a healthy donor was obtained from the local blood donation center under informed consent. The buffy coat was processed as described previously [25]. The buffy coat was diluted to 180 mL in 0.6 % (w/v) sodium citrate in PBS adjusted to pH 7.2 at 4 °C (SC buffer), carefully layered onto Lymphoprep™ iso-osmotic medium and centrifuged at 800 × *g* for 30 min without brake and with minimal acceleration at RT. Mononuclear cells were washed 5 × with SC buffer to remove all Lymphoprep™, and cryogenically stored in liquid nitrogen until further use. Upon use, cells were thawed and used without passaging. A purified monocyte fraction was isolated from the thawed cells using the negative selection MACS^®^ Pan Monocyte Isolation Kit (Miltenyi Biotec) with LS columns according to the manufacturers’ instructions. After isolation, cells were resuspended in Neutral medium (Table 1). This purified monocyte fraction will from now on be referred to as ‘monocytes’.

**Table 1:** Media names, components and functions within the context of this study.

| Name | Components | Function |
| --- | --- | --- |
| OB medium | DMEM low glucose<br>10 % FBS<br>1 % anti-anti<br>50 µg/mL L-ascorbic-acid-2-phosphate<br>100 nM dexamethasone<br>10 mM β-glycerophosphate | Osteogenic medium. One of three media compared during the co-culture. This medium was expected to further stimulate mineralized matrix deposition by OB. |
| Neutral medium | RPMI-1640<br>10 % FBS<br>1 % Anti-Anti | Unsupplemented medium. One of three media compared during the co-culture. This medium was expected to allow OB and OC crosstalk to control ongoing matrix deposition and resorption. |
| OC medium | RPMI-1640<br>10 % FBS<br>1 % Anti-Anti<br>50 ng/mL M-CSF<br>50 ng/mL RANKL | Osteoclastogenic medium. One of three media compared during the co-culture. This medium was expected to stimulate OC resorption. |
| MSC expansion medium | DMEM high glucose<br>10 % FBS<br>1 % anti-anti<br>1 % NEAA<br>1 ng/L bFGF | Used to expand MSCs prior to seeding onto scaffolds. |
| MSC seeding medium | DMEM high glucose<br>10 % FBS<br>1 % anti-anti | Unsupplemented medium that is used for MSC seeding onto scaffolds and prewetting of scaffolds. |
| Monocyte priming medium | RPMI-1640<br>10 % FBS<br>1 % Anti-Anti<br>50 ng/mL M-CSF | Used to prime monocytes during the first two days of culture which benefits osteoclastogenic differentiation. |

#### 2D OC culture and analysis

To verify that the monocytes can form multinucleated TRAP expressing and resorbing OC-like cells, 0.25 × 10^6^ monocytes per cm^2^ (n = 4 per group) were seeded in monocyte priming medium (Table 1) on 24-well Corning^®^ osteo assay plates and regular tissue culture plastic 24-well tissue culture plates in monolayer. Monocyte priming medium was replaced with OC medium (Table 1) or Neutral medium after 48 h. Medium was replaced 3 × per week for 14 d. The Corning^®^ osteo assay plate from the 2D OC culture was analyzed for resorption according to the manufacturers’ instructions. Cells were removed using 5 % bleach for 5 min. The plate was washed with UPW and dried at 50 °C. Bright field images were taken with a Zeiss Axio Observer Z1 microscope and binarized with Matlab^®^. The 2D OC culture in plastic well-plates was immunofluorescently labelled for OC markers (TRAP or Integrin β3), actin (TRITC-conjugated-Phalloidin) and nuclei (DAPI). Fluorescence images were taken with a Zeiss Axiovert 200M microscope. Supernatant culture medium samples were taken and stored at −80 °C at each medium change and analyzed for TRAP enzyme activity as described later for the 3D co-culture.

#### Fabrication of silk fibroin scaffolds

Silk fibroin (SF) scaffolds were produced as previously described [24,25,29,30]. Unless stated otherwise, solutions used were ultra-pure water (UPW) or dissolved in UPW. Cocoons from the *Bombyx mori L*. silkworm were degummed by boiling in 0.2 M Na_2_CO_3_ twice for 1 h, rinsed (boiling UPW) followed by 10 × washing (cold UPW). After overnight drying the silk was dissolved in 9 M LiBr (10 % w/v) at 55 °C for 1 h and filtered through a 5 μm filter after cooling to RT. The filtered silk solution was dialyzed for 36 h using SnakeSkin Dialysis Tubing. UPW was refreshed at 1, 3, 12, and 24 h. The dialyzed solution was frozen (−80 °C), lyophilized and dissolved to a 17 % (w/v) solution in 1,1,1,3,3,3-Hexafluoro-2-propanol (HFIP). 1 mL silk-HFIP was added to 2.5 g NaCl (granule size between 250-300 μm) in a Teflon container and was allowed to dry at RT for at least 4 d. β-sheet formation [31] was induced by immersion in 90 % (v/v) methanol for 30 min. Silk-salt blocks were dried at RT overnight and cut into 3 mm discs using a precision cut-off machine (Accutom-5^®^, Struers GmbH Nederland, Maassluis, the Netherlands). NaCl was leached out in UPW which was refreshed after 2, 12, 24 and 36 h. Scaffolds were punched with a biopsy punch (5 mm) and sterilized by autoclaving (20 min at 121 °C) in phosphate buffered saline (PBS). Scaffolds were pre-wetted in mesenchymal stromal cell (MSC) seeding medium (Table 1) prior to use.

#### Construct mineralization by hMSC-derived OBs

The human hMSCs used in this study were previously isolated from human bone marrow and characterized [32] and were used as described previously [25]. Briefly, 2.5 × 10^3^ cells/cm^2^ (passage 5) were seeded and expanded for 6 d in MSC expansion medium (Table 1). 1 × 10^6^ hMSCs in 20 μL MSC seeding medium were seeded onto each pre-wetted scaffold and incubated for 90 min at 37 °C and are from now on referred to as constructs. These constructs were then transferred to 8 custom-made spinner flask bioreactors (n = 4 per bioreactor) containing magnetic stir bars as described previously [24,25] that were filled with 5 mL OB medium (Table 1). Each bioreactor was placed in an incubator (37 °C, 5 % CO_2_) on a magnetic stirrer plate (RTv5, IKA, Germany) rotating at 300 RPM [24]. Medium was changed 3 times a week for 11 weeks. The resulting cells are from now on referred to as OBs.

#### Initiation of 3D co-culture on pre-mineralized constructs

Constructs which had been in culture for 11 weeks with osteogenically stimulated hMSCs (from now on referred to as OBs) were incised with a 4 mm deep incision in the transverse plane to allow seeding to the center of the constructs. 1 million (M) monocytes, 1.5 M monocytes or no monocytes in 7.5 μL monocyte priming medium (Table 1) were seeded into the incision of constructs pre-wetted in monocyte priming medium. All constructs were incubated for 180 min at 37 °C to facilitate cell attachment. Then, all constructs were placed back into the bioreactors (n = 4 per bioreactor) with 5 mL monocyte priming medium per bioreactor. No stirring was applied during the co-culture to better stimulate monocyte attachment and differentiation [33–35]. The 3D co-culture of OBs with monocytes will be referred to as ‘3D co-culture’. The 3D co-cultures were primed in monocyte priming medium for 48 h [36,37]. Monocyte priming medium was replaced after 48 h with one of three media for the remainder of the culture: OB medium, Neutral medium, or OC medium (Table 1). This resulted in bioreactors with constructs seeded with 0 M monocytes cultured in OB and OC medium, 1 M monocytes cultured in OB, OC and Neutral medium, and 1.5 M monocytes cultured in OB, OC and Neutral medium. Medium was replaced 3 × per week for 21 d.

#### μCT imaging

μCT measurements were performed as previously described [25] on a μCT100 imaging system (Scanco Medical, Brüttisellen, Switzerland) every week except week 2 of the 3D OB mono-culture to monitor tissue mineralization. After 11 weeks, co-culture was initiated and the scanning frequency was increased to twice per week (isotropic nominal resolution:17.2 μm, energy level: 45 kVp, intensity: 200 μA, integration time: 300 ms, two-fold frame averaging, computed tomography dose index (CTDI) in air: 230 mGy). A constrained Gaussian filter was applied to reduce part of the noise (Filter support: 1.0, filter width sigma: 0.8 voxel). A fixed region of interest (RoI) of 205 slices was selected within each bioreactor. This ensured that the same RoI of every scaffold was scanned each time and limited the required scan time and radiation exposure to 30 min per scan. At the start of the co-culture, this RoI was reassessed for each bioreactor to contain as much of the constructs as possible, and this exact RoI of 205 slices was used for the remainder of the co-culture. Segmentation was done at a global threshold of 23 % of the max greyscale value. Image processing language (IPLFE v2.03, Scanco Medical AG) was used to further process the images. Component labelling was used to remove unconnected objects < 50 voxels.

These were neglected from further analysis. The mineralized tissue volume of the RoI was assessed using quantitative morphometry. 3D OB mono-culture (mineralized construct generation) quantitative μCT data was used as measured, whereas 3D co-culture quantitative μCT data was normalized to the mineralized volumes of the first scan of each individual construct during the co-culture (d 4 of co-culture) counting that volume as 100 %. All successive scans were presented as volume change with respect to this first scan. Rigid 3D registration was used to register the follow-up (d 7) to the baseline image (d 4) of the 3D co-culture [38]. Color coding was used to label resorption (blue), formation (orange) and unaltered regions (grey). Unconnected objects < 100 voxels were removed as before using component labelling for registered images only.

#### Histology

At the end of the culture, constructs were fixed in 10 % neutral buffered formalin for 24 h at 4 °C. Fixed constructs were dehydrated with an EtOH and xylene series (1.5-2 h per step) and embedded in paraffin. 10 μm thick vertical sections were mounted on poly-L-lysine coated microscope slides. Sections were dewaxed and rehydrated with a Xylene and EtOH to UPW series. These sections were used for histology and immunofluorescence. Sections were stained with von Kossa staining to visualize calcium phosphate deposition (30 min in 1 % aqueous silver nitrate (w/v) under UV light, rinsed with UPW, 5 min in 5 % sodium thiosulfate (w/v), rinsed with UPW, 5 min in nuclear fast red, rinsed in UPW). To visualize calcium deposition, sections were stained for Alizarin Red (2 min in 2 % Alizarin Red in H_2_O, pH 4.2). Stained sections were dehydrated using EtOH (von Kossa) or acetone (Alizarin Red) to Xylene and coverslipped with Entallan^®^ Bright field images were taken with a Zeiss Axio Observer Z1 microscope.

#### Immunofluorescence

Sections were prepared, dewaxed, and rehydrated as for histology. After antigen retrieval (citrate buffer at 95 °C for 20 min, then slowly cooled back to RT), cross-reactivity was blocked (10 % donkey serum for 30 min). Primary antibodies were incubated at 4 °C overnight, and secondary antibodies were incubated at RT for 1 h. Sections were labelled for OC marker (integrin β3) [39,40], actin (TRITC-conjugated-Phalloidin) and nuclei (DAPI). Sections were coverslipped with Mowiol^®^ and imaged with a Leica TCS SP5X microscope.

#### Scanning electron microscopy (SEM)

Constructs for SEM were fixed using glutaraldehyde (2,5 % for 24 h at 4 °C), dehydrated with a graded EtOH series followed by a graded 1,1,1-Trimethyl-N-(trimethylsilyl)silanamine (HMDS)/ethanol series, dried overnight at RT and sputter coated with 5 nm gold (Q300TD, Quorum Technologies Ltd, Laughton, UK). Sputter coated constructs were imaged with SEM (Quanta600, FEI Company, Eindhoven, the Netherlands, spot size 3.0, 5.00 kV, working distance 10 mm).

#### Tartrate-resistant acid phosphatase (TRAP) quantification in supernatant

Supernatant medium samples were taken and stored at −80 °C just before each medium change (n = 4 technical replicates per bioreactor). 20 μL of the supernatant medium samples or nitrophenol standard in PBS were incubated in translucent 96-well plates at 37 °C for 90 min with 100 μL para-nitrophenylphosphate (pNPP) buffer (1 mg/mL pNPP, 0.1 M sodium acetate, 0.1 % (v/v) triton-X-100 in PBS adjusted to pH 5.5 supplemented with 30 μl/mL tartrate solution). The reaction was stopped with 100 μl 0.3 M NaOH. TRAP enzyme activity was determined by measuring absorbance at 405 nm and recalculated to pNPP transformation per minute.

#### Statistical analysis

Quantitative data is represented as mean ± standard deviation (SD) and was analyzed using GraphPad Prism version 8. Data used for statistical analysis was tested for normality using the Shapiro-Wilk normality test and was normally distributed. Groups were compared using a Two-Way Analysis of Variances (ANOVA). Trends within groups over time were compared using a Repeated Measures ANOVA. Planned comparisons within groups were: volume increase within the 1^st^ week of co-culture, volume decrease starting after the 1^st^ week, TRAP release increase starting in the 1^st^ week, TRAP release decrease starting after the 2^nd^ week. Bonferroni correction was used to account for multiple comparisons in all other comparisons. Geisser-Greenhouse correction was used to account for unequal variances. Differences were considered statistically significant at a level of *p* < 0.05. SDs that were much larger than others within the same dataset were tested with Grubbs test for outliers against other SDs within the dataset. If an SD was positively identified, the underlying data was searched for outliers. Statistical analyses were rerun with the identified datapoint replaced with the mean of the remaining datapoints. If this led to different significances, then figures show a ‘+’ indicating a relevant outlier. The original (unchanged) dataset is shown in all figures regardless of the outcome, but the results of both analyses are described in the results section. Grubbs test was used for the TRAP results of 1 M monocytes seeded in both OC medium and Neutral medium, timepoints of d 7 and d 16 respectively. Notable significant effects are numbered in the results section and in the figures using unique sequential numbering preceded by an asterisk throughout the study for easy referencing between texts and figures.

## Results

### Verification of osteoclastogenesis in 2D

Monocytes were cultured in 2D to verify their capability to differentiate into multinucleated TRAP expressing resorbing cells. Monocytes cultured in OC medium continuously released increasing amounts of TRAP into the culture supernatant over a period of 14 d ending at 2.84 ± 0.29 μmol/min, whereas those cultured in Neutral medium released lower quantities of TRAP even at the peak of 1.37 ± 0.08 μmol/min at d 7 (difference with OC group at d 7: 0.54 μmol/min, *p* = 0.0089, *1) after which the release of TRAP decreased again (Fig 1A). After 14 d on Osteo Assay plates, cells cultured in OC medium showed extensive resorption, whereas those cultured in Neutral medium showed only minimal traces of resorption (Fig 1B + C). Fluorescence imaging revealed that cells cultured in OC medium developed into large multinucleated cells with clearly defined actin rings, expressing OC markers TRAP (Fig 1D) and integrin-β3 (Fig 1F), although TRAP was also expressed in unfused monocytes. In Neutral medium some clusters of nuclei are seen, possibly small multinucleated cells that developed as a result of 2-day priming (Fig 1E). Most monocytes in the Neutral medium group expressed TRAP like those in the OC medium group, whereas integrin-β3 was expressed almost exclusively in multinucleated cells and was not found in cells cultured in Neutral medium (Fig 1G). These results confirm that the monocytes used in this study were able to differentiate into multinucleated TRAP expressing and resorbing OCs in 2D.

**Fig 1.**
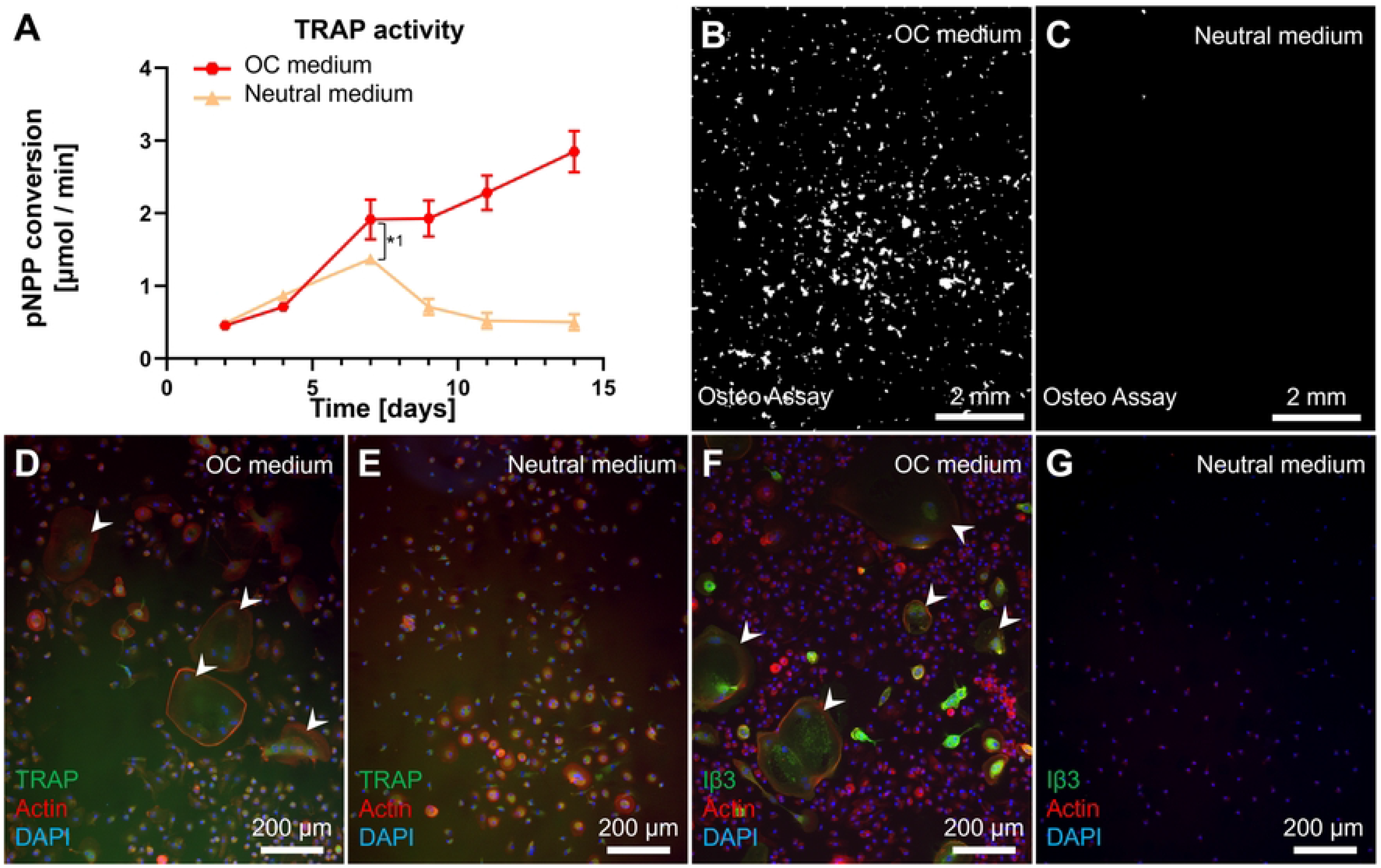
Monocytes can differentiate into TRAP expressing resorbing cells in 2D mono-culture. **(A)** TRAP expression when monocytes are cultured in OC medium or Neutral medium. Both groups showed an initial increase of TRAP activity, but only the osteoclastogenically stimulated group kept releasing more TRAP after d 7. **(B)** Resorption of Osteo Assay plate surfaces after 14 d of culture in OC medium or **(C)** Neutral medium. Images are binarized light microscopy images of the center of the well. Resorption was present in the OC medium group, whereas resorption in the Neutral medium group was negligible. **(D)** Multinucleated (blue) TRAP (green) expressing cells (white arrowheads) with a clearly defined actin ring (red) were seen when cultured with OC medium, whereas monocytes cultured in Neutral medium **(E)** mostly remain uninuclear and only lightly express TRAP. **(F)** Integrin β3 (green) was expressed almost exclusively in multinucleated cells (white arrowhead) in a culture with OC medium, whereas monocytes cultured in Neutral medium **(G)** did not express Integrin β3 at all.

### Mineralized matrix is deposited onto SF scaffolds

hMSCs were seeded onto SF scaffolds (Fig 2A + B) and differentiated into mineralized matrix depositing MSC-derived OBs for 11 weeks. Matrix deposition was monitored using μCT for each individual construct until the start of co-culture (Fig 2C + D). Non-mineralized SF scaffolds were not detectable with the used μCT settings. Already after 6 d, 0.002 ± 0.003 mm^3^ of mineralized matrix was detected with μCT. Mineralized matrix deposition continued steadily throughout the culture duration and throughout the construct reaching a mean mineralized volume of 9.67 ± 2.42 mm^3^ on d 69.

**Fig 2.**
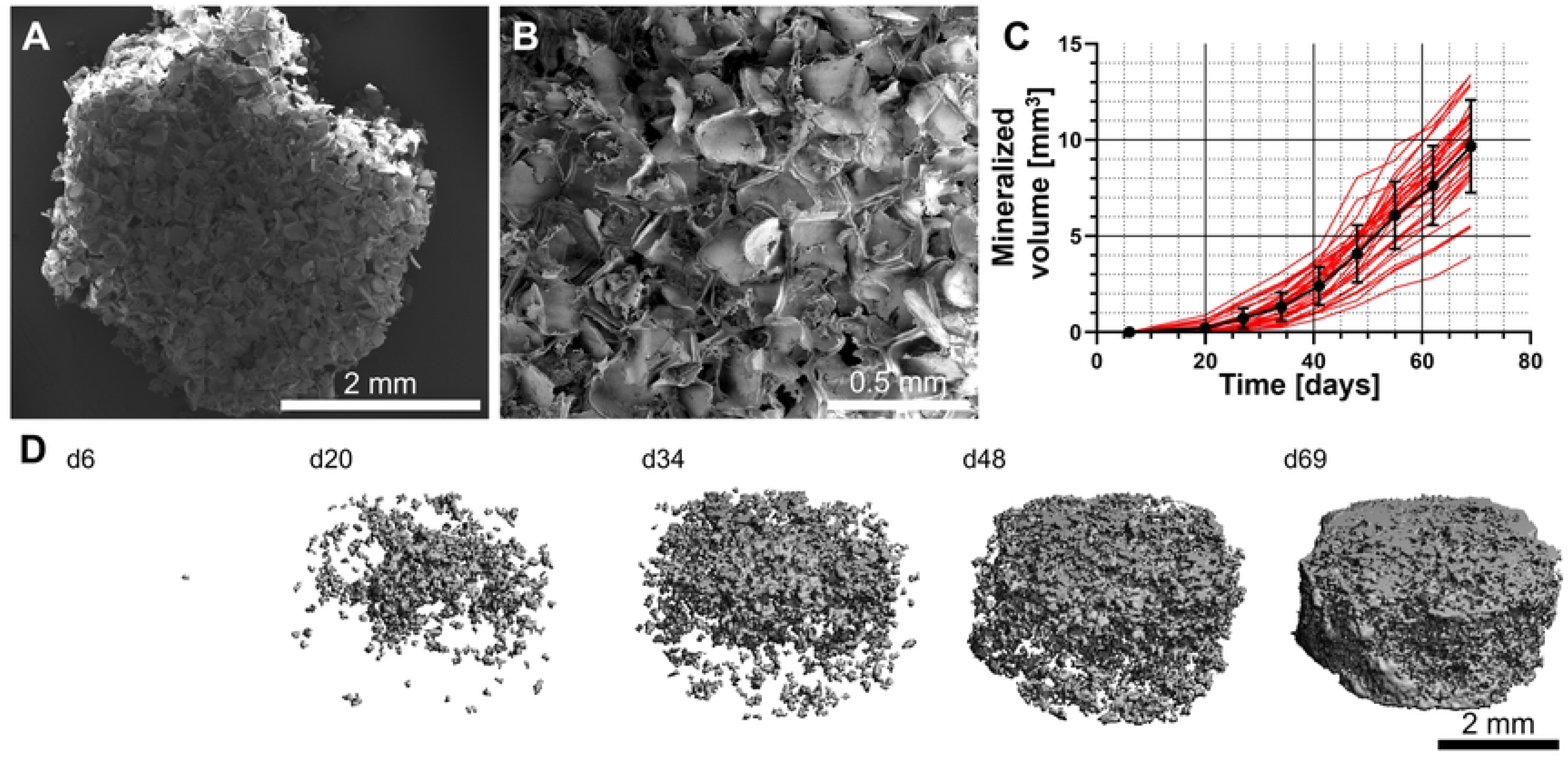
Mineralized matrix deposition over time onto SF scaffolds. **(A)** Top view of a freshly prepared SF scaffold. **(B)** Higher magnification view revealing the highly porous nature of the scaffolds. **(C)** μCT monitoring confirmed continuous mineralized matrix deposition and an increase in mineralized volume during the entire culture duration. Individual construct volumes are shown in red; the mean ± SD are shown in black. **(D)** Representative images of one construct over time showing the volumetric distribution and growth of an individual construct.

### Histology and SEM show calcium phosphates and ECM presence after co-culture

An incision in the transverse plane (Fig 3A) was used to deliver cells to the center of the mineralized construct. This seeding strategy was chosen because otherwise, the deposited (mineralized) tissue (Fig 3B) could have prevented the subsequently seeded monocytes from penetrating deeper into the construct if seeded on the outside surface. Alizarin Red (Fig 3C) and von Kossa (Fig 3D) stainings showed that calcium phosphate deposits are abundantly present in the ECM in all groups at the end of co-culture.

**Fig 3.**
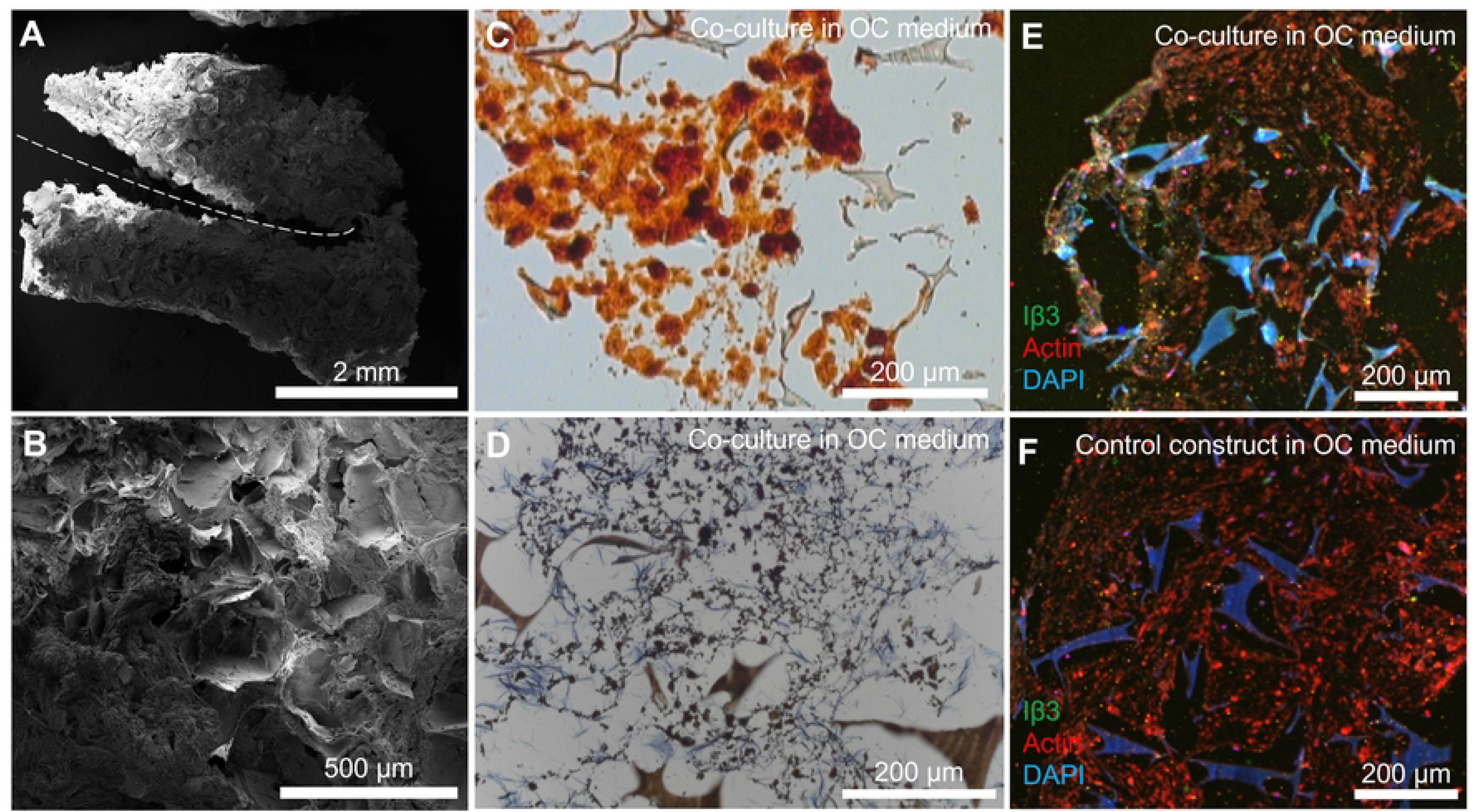
Construct morphology and histology after co-culture. **(A)** Mineralized constructs were sectioned in the transverse plane to seed the monocytes. The incision is marked with a dashed line. **(B)** After co-culture, ECM deposition into the pores of the construct was visible on SEM images. **(C)** Alizarin Red staining confirmed the presence of calcium throughout the constructs after co-culture in OC medium. **(D)** Von Kossa staining confirmed the deposition of (calcium) phosphates throughout the constructs after co-culture in OC medium. **(E)** Immunofluorescence for Integrin β3 (green), actin (red) and nuclei (blue) revealed the presence of OC marker Integrin β3 even after 21 d of co-culture in constructs cultured in OC medium. **(F)** Immunofluorescence for Integrin β3 (green), actin (red) and nuclei (blue) on a control construct without monocytes, cultured in OC medium.

### Confirmation of osteoclastogenesis on mineralized constructs

Immunofluorescence imaging revealed the presence of OC marker integrin-β3 in constructs cultured in OC medium even after 3 weeks of co-culture while the marker was absent in constructs without seeded monocytes (Fig 3E + F). SEM images were taken after 3 weeks of co-culture to investigate the presence of monocytes and OCs (Fig 4). Only in the groups cultured in OC medium, large OC-like cells were identified (Fig 4A + B), sometimes in the presence of what could be resorption trails (Fig 4C). Small round monocyte-like cells were also identified in all groups in which monocytes were seeded (Fig 4D + E). MSC-produced extracellular matrix was present abundantly throughout all constructs, often completely filling pores in the SF scaffolds (Fig 4F).

**Fig 4.**
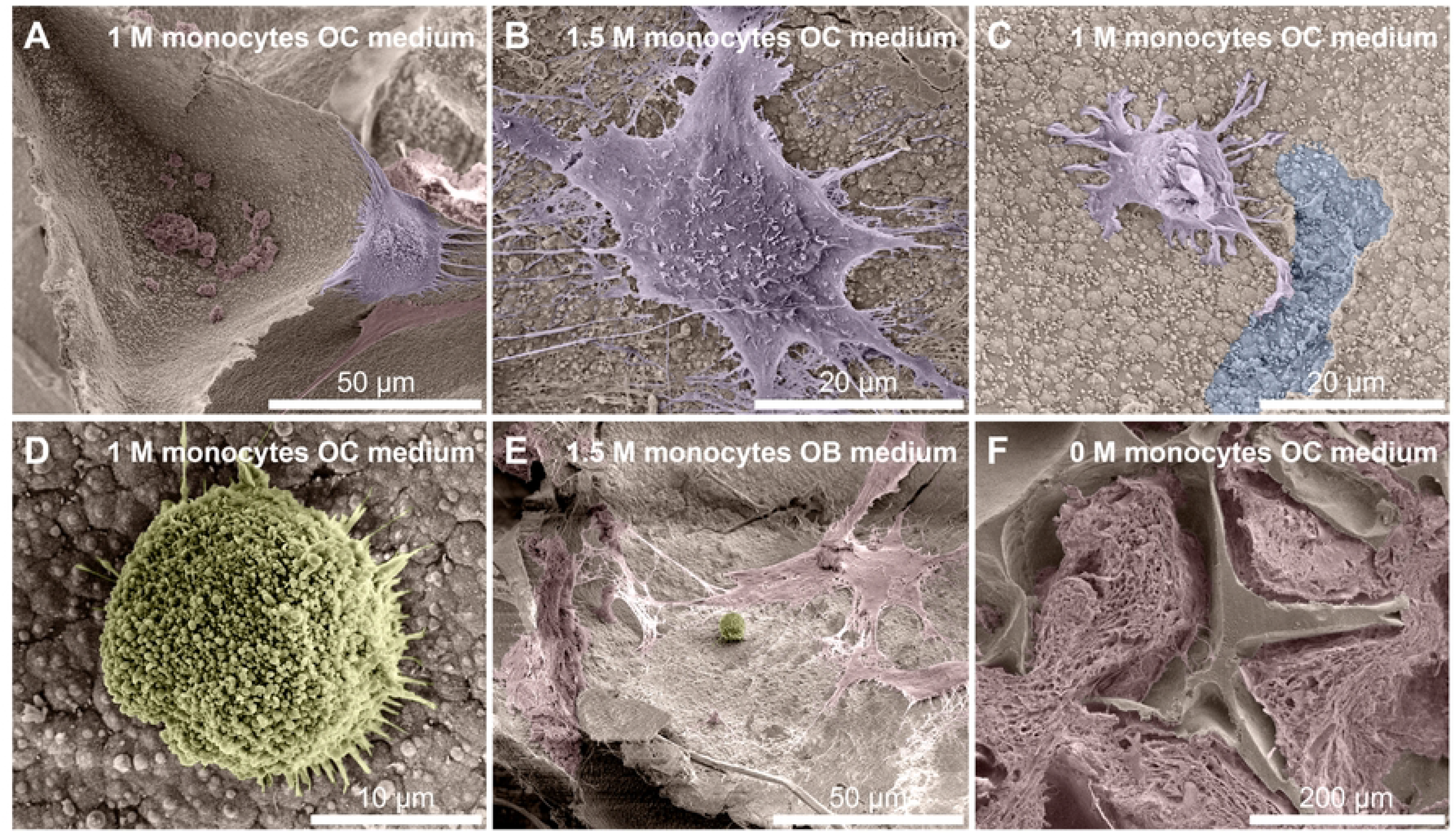
Cells and tissue after co-culture. **(A)** An OC on the edges of a pore, with likely remnants of MSC-derived cells and ECM at the bottom of the pore. **(B)** OC with many filopodia stretching out in all directions. **(C)** Small OC-like cell moving away from a resorption trail. Its OC lineage is recognizable by the ‘frizzled’ appearance on the top surface like that of the OC in figs 4A and 4B, which contrasts with the smoother surface of MSC-derived cells and ECM that is marked in red in for example fig 4A and 4E. **(D)** Close-up of a monocyte. Round cells such as these were only found in constructs onto which monocytes were seeded and not on unseeded control constructs, regardless of media type. **(E)** A lone monocyte amidst deposited MSC-derived cells and/or ECM. **(F)** Overview image of the native SF scaffold structure that has been filled with ECM. Images are digitally enhanced SEM images. OC are colored purple, monocytes are colored yellow, resorption trails are colored blue, MSC/OB derived ECM (possibly including cells) is colored red. Original unenhanced images are available in S1_Figure_unenhanced.

### Co-culture media affects the amount TRAP release

Monocytes were co-cultured with OBs on constructs in one of three media: OC medium, Neutral medium, or OB medium. As expected, TRAP release (Fig 5A) over time was highest in the group cultured in OC medium (1.58 ± 0.20 μmol/min at d 14). TRAP release in the OC medium group was significantly higher than in the Neutral medium group from d 7 to d 18 (*p* < 0.05 for all timepoints, d 7 only after outlier correction, *3). TRAP release in the Neutral medium group increased compared to the group cultured to OB medium only after 11 d (*p* = 0.04 at d 14, *p* = 0.02 at d 16 after outlier correction, *p* = 0.99 with outlier, *2), but TRAP release in the Neutral medium group followed a clear linear trend (*p* = 0.0185). This suggested that differentiation towards OCs and onset of increased TRAP release occurred later in the Neutral medium group than in the OC medium group. Remarkably, monocytes cultured in unfavorable OB medium still released TRAP into the medium, as it is structurally higher than the ‘baseline’ TRAP measurement of constructs onto which no monocytes were seeded (*p* < 0.05 at each timepoint).

**Fig 5.**
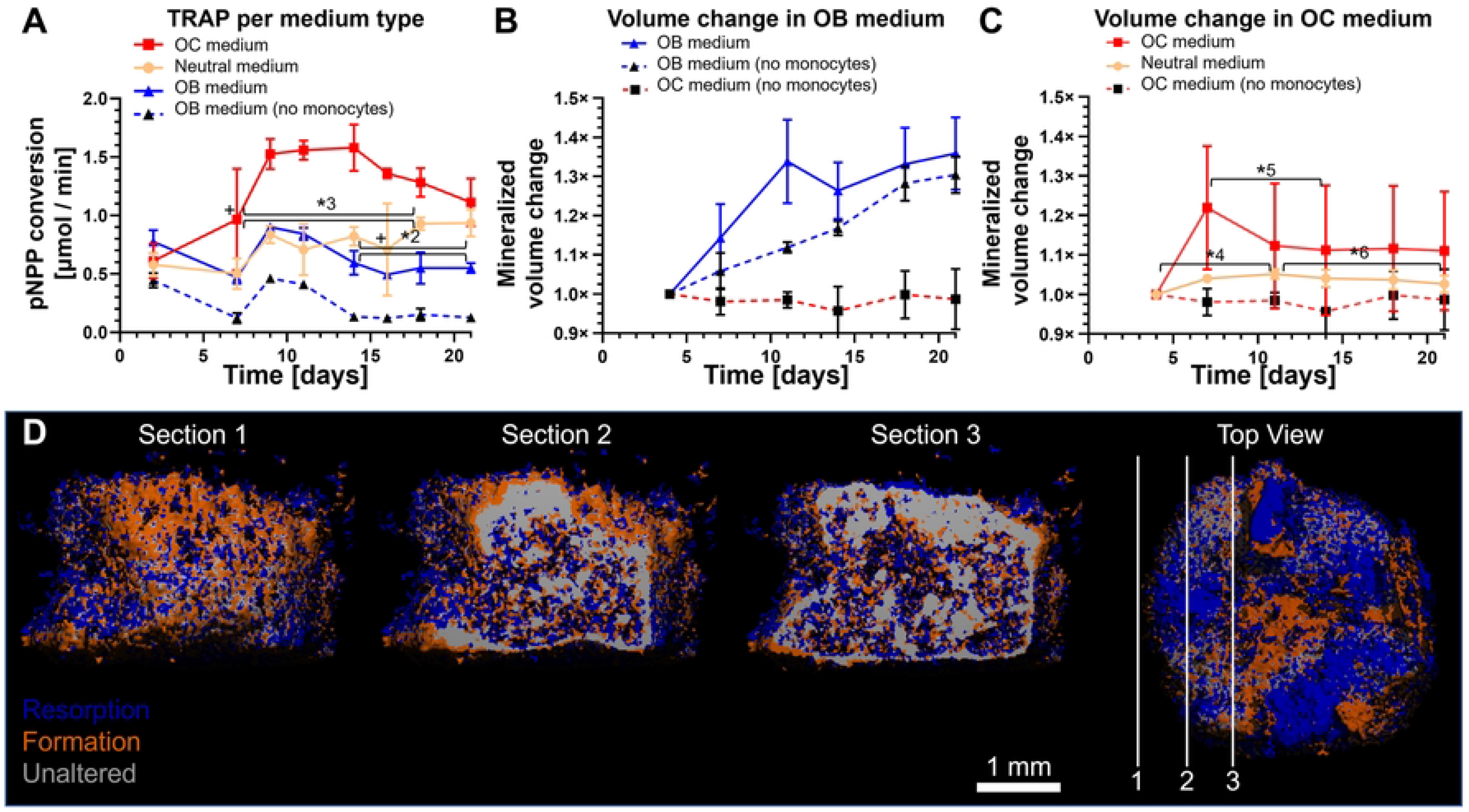
Medium composition affects TRAP activity, mineralized tissue formation and resorption activity in co-culture in 1 M seeded monocytes groups. **(A)** TRAP release of the 1 M monocytes groups. Co-cultures in OC medium showed highest TRAP release, followed by those cultured using Neutral medium and OB medium. The 0 M group in OB medium serves as reference. **(B)** Mineralized volume increased in OB medium with or without co-culture, but no longer increased in OC medium without monocytes. **(C)** Mineralized volume change of constructs of co-culture in OC or Neutral medium. There was no resorption without monocytes. With monocytes, mineralized volume increased in the first few days, and then decreased. The OC medium group showed a large and early decrease in volume, while the Neutral medium group showed a small but steady decrease in volume until the end of culture. The OC medium (no monocytes) group is identical in panels B and C and is for reference. **(D)** Three sections of the same registered scans of d 4 and d 7 of co-culture construct cultured in OC medium show many resorption (blue) and formation (orange) events on the pore surfaces, while the inside of the SF structure of the construct remains mostly unchanged (grey). The locations of the sections within the construct are shown on the top view.

### Co-culture medium affects mineralized matrix volume

Mineralized volume was measured over time with μCT and normalized relative to the measurement at d 4 of co-culture. As expected from the mineralization curve before co-culture (Fig 2C), the amount of mineralized volume in constructs cultured in OB medium both with and without seeded monocytes increased for another 3 weeks (Fig 5B). With exception of d 11 (*p* = 0.025), the volume change per timepoint between these groups were not significantly different suggesting that the mere presence of monocytes did not influence OB activity. Switching from OB medium to OC medium or Neutral medium seemingly ended this trend regardless of monocyte presence (Fig 5C). The presence of monocytes seemed to prolong mineralized matrix deposition by a few days, but this effect was only statistically significant for the co-culture in Neutral medium (*p* = 0.0035, *4) and not for the co-culture in OC medium (*p* = 0.064), likely because of the large SDs withing this group. In all individual constructs of the OC medium group, there was a significant decrease in mineralized volume between d 7 and d 14 (9.12 % decrease, *p* = 0.003, *5), suggesting differentiation and peak OC resorption activity happened in this period. In the Neutral medium group, a smaller but gradual decrease in mineralized volume was seen between d 11 and d 21 (2.5 % decrease, p = 0.018, *6). Formation and resorption occurred both on the inside and the outside of the constructs. An example μCT image registration shows the difference between d 4 and d 7 in a co-culture in OC medium (Fig 5D).

### Monocyte seeding density affects TRAP release but not net resorptive activity

Monocytes were seeded at a density of either 1 M or 1.5 M cells per construct. A higher seeding density led to a seemingly higher TRAP release (Fig 6A + C), but these differences were only significant in the group cultured in Neutral medium (Fig 6C) (*p* < 0.05 for all timepoints, d 16 only after outlier correction, *7) suggesting that the presence of more neighboring monocytes contributed to differentiation by giving more chances for cell fusion and subsequent TRAP release when no OC supplements were present. An excess of OC factors (Fig 6A) partially bypassed the need to have many neighboring cells as it resulted in higher TRAP expression in the group seeded with only 1 M monocytes.

**Fig 6.**
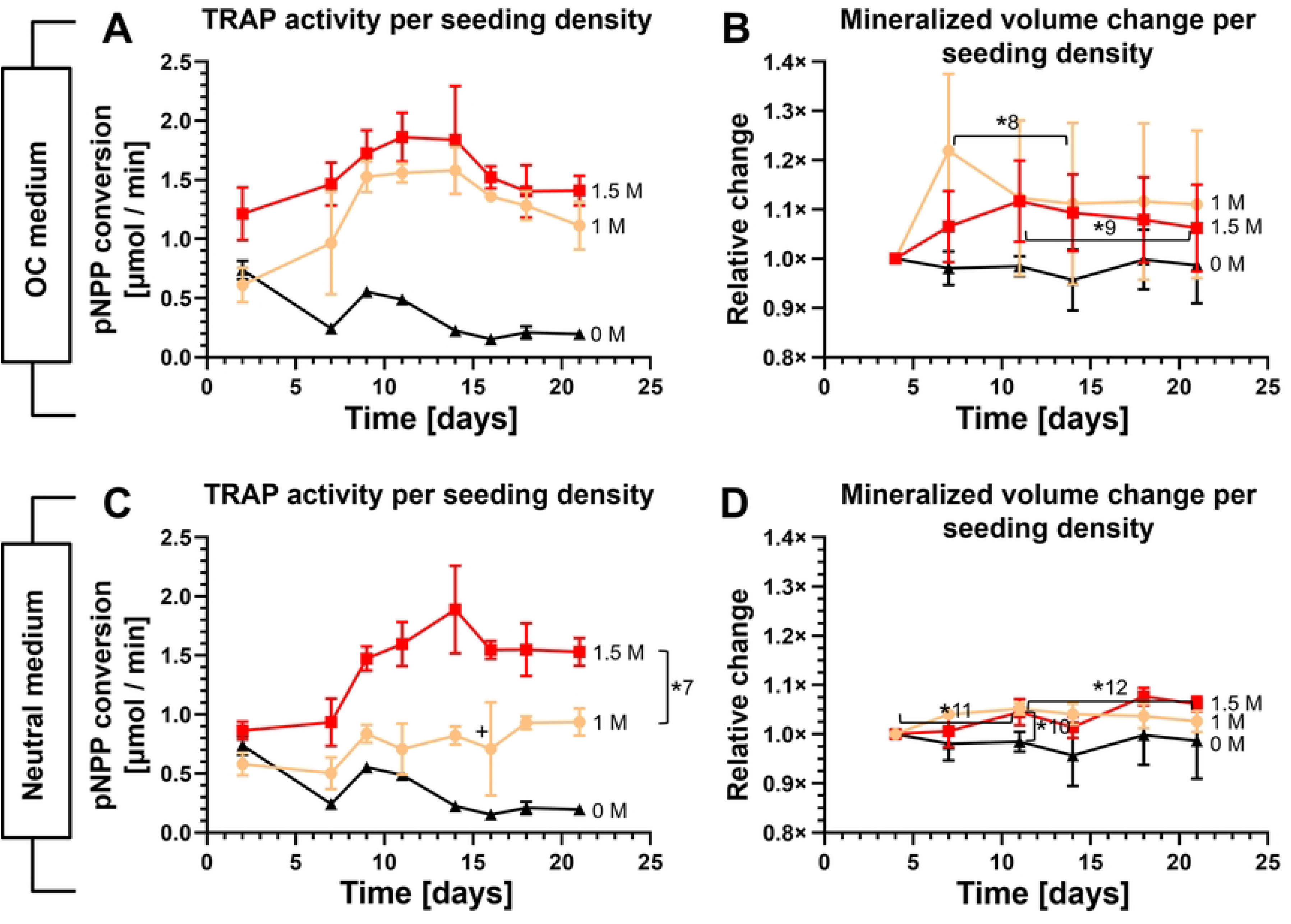
TRAP release and resorption during co-culture at different seeding densities in different media. **(A)** TRAP release of all monocyte seeding densities cultured in OC medium. TRAP expression was higher in constructs with 1.5 M than in those with 1 M monocytes seeded, but not significantly. **(B)** Mineralized volume change of all monocyte seeding densities cultured in OC medium. The control constructs showed no net change in mineralized volume. Both monocyte seeding densities increased in mineralized volume, followed by a downward trend in mineralized volume. **(C)** TRAP release of all monocyte seeding densities cultured in Neutral medium. The co-cultures with 1.5 M monocytes showed more TRAP release than the co-cultures with 1 M monocytes. **(D)** Mineralized volume change in co-cultures cultured in Neutral medium. The mineralized volume of the 1 M and 1.5 M groups increased significantly until d 11. The 1 M group steadily decreased in volume from d 11 to the end of co-culture. Note that the 0 M and 1 M lines from all panels of this figure were also shown in Fig 5. The unseeded controls in Fig 6a and b were cultured in OC medium and are show as a reference in these figures only.

In line with the TRAP results, the effects of seeding density did not result in significant differences in resorption between the groups cultured in OC medium (*p* > 0.05 at each timepoint) (Fig 6B), although there was a slight downward trend in mineralized volume (indicating resorption) in both the 1 M monocytes (d 7 to d 11, *p* = 0.008, *8) and the 1.5 M monocytes (d 11 to d 21, *p* = 0.019, *9) group. This was not seen in the control group. In the groups cultured in Neutral medium (Fig 6D), mineralized volume in the 1 M and 1.5 M monocytes group was significantly different from the control group without monocytes only at d 11 (1 M: *p* = 0.0023, 1.5 M: *p* = 0.0123, *10), around the time where OC are expected to become active as shown by TRAP results. Other than this there were no significant differences between groups. Both the 1 M monocytes group (5.2 % increase, *p* = 0.0035, *11) and the 1.5 M monocytes group (4.5 % increase, *p* = 0.043, *11) slightly but significantly increased in mineralized volume during the first 11 d of co-culture suggesting again that monocytes briefly stimulate mineralization. The constructs of the 1 M monocytes group slightly but continuously decrease in volume from d 11 until d 21 (*p* = 0.018, *12) as shown earlier in Fig 5D.

## Discussion

If 3D OB-OC co-culture models are to be used for fundamental research, drug development or personalized medicine, it is imperative that these models can demonstrate (im)balanced matrix formation and resorption. Our earlier work has shown that both formation and resorption can be monitored within 3D constructs in co-culture [25]. Resorption occurred in the presence of OC supplements, which are potent and potentially overshadowed any cell-cell communication, enforcing a state of maximum resorption. For fundamental research and drug testing, the cells should appropriately respond to the introduced stimuli in a concerted reaction. This is only possible if supplements don’t overrule the cell response. Ideally, the model allows to investigate remodeling both in equilibrium or out of balance situations, mimicking a healthy state or a diseased state, respectively. It has previously been shown that both medium composition [26,27] and cell seeding density [28] affect intercellular communication, maturation, differentiation and cell activity. In this study, different media compositions and seeding densities were used to tune the direction and amount of remodeling to a state of equilibrium, forced formation and forced resorption.

Co-culture in OB medium resulted in continued mineralized matrix deposition. Mechanical loading was used for its stimulatory effect on mineralized matrix deposition [24,35,41] during construct mineralization before the co-culture. Mechanical loading was disabled during the co-culture to facilitate undisturbed monocyte attachment and differentiation. The continued matrix deposition suggests that, while mechanical stimulation is beneficial for matrix deposition, it may not be necessary to have mechanical loading once mineralization is ongoing. The presence of monocytes did not significantly affect mineral deposition. This was expected because the co-culture of this group was provided with an excess of osteogenic supplements, effectively overruling OC signaling that could affect mineralized matrix deposition. Another factor that could have contributed to mineralization was the choice of FBS, which has been shown to contain intrinsic alkaline phosphatase activity, the ability to cause calcium phosphate deposition even in the absence of cells, and greatly affect osteogenic differentiation [42]. At the same time, the TRAP measurements suggested that there was only a ‘basal’ expression of TRAP by monocytes [25] and minimal OC differentiation ongoing [43,44]. This was in line with what was expected based on the bone remodeling cycle, where a resorption phase precedes a reversal phase [45] followed by a formation phase to repair the resorbed area. If OB stimulation simulates the formation phase, there would be no direct need for a new resorption phase, and OBs would not want to stimulate additional osteoclastogenesis. *In vivo*, this task is attributed to the osteocytes [1,46]. While the current study did not attempt to prove their presence, earlier work has shown indications of osteocyte-like cells in similar culture conditions [25]. The model with OB medium could be used to investigate the maximum mineralizing capacity of different cell donors (such as osteoporotic patients) within the context of the model.

Monocyte presence prolonged OB mineralized matrix deposition. In both Neutral and OC medium, when monocytes were present, there was ongoing mineralized matrix deposition in the first 11 d. Compared to OB medium however, the mineralization curve was flattened. This curve was absent in the group where no monocytes were seeded. This indicated that in the absence of osteogenic supplements, the presence of monocytes was able to marginally support further mineralization. The net effect of this activity dissipated after d 11, as monocytes started to differentiate into OCs and became capable of resorbing matrix, which likely resulted in formation and resorption masking each other’s changes in mineralized volume. This was further demonstrated by the μCT registrations revealing both resorption and formation occurring simultaneously. This confirmed that the model using Neutral or OC medium can be used to monitor both resorption and formation in co-culture simultaneously.

In OC medium, resorption initiated faster than in Neutral medium. The OC medium group had a large increase in resorption during week 2 of co-culture while the mineralized volume in Neutral medium decreased much slower over the remaining culture period. This is in line with the TRAP release results that showed a similar peak for the OC medium during week 2 of co-culture while the Neutral medium group gradually released increasing amounts of TRAP over time. The early increased TRAP release and faster onset of resorption in the OC medium group likely resulted from more OCs generated by an excess of osteoclastic supplements, which is in line with the identification of OC-like cells and resorption trails in SEM images especially in this group [47–49]. Cell behavior in the Neutral medium group relied solely on the interaction with the matrix and on OB-OC crosstalk since no OC supplements supporting OC differentiation were added. Nevertheless, a slow build-up of TRAP release and by extension OC differentiation took place, although it did not reach the same level as in the OC medium group. This confirmed that when OC medium is used, monocytes are forcefully steered towards osteoclastogenesis, whereas Neutral medium allows exclusively OB-OC crosstalk and basic medium components [42] to regulate remodeling. The model using OC medium would be suitable to study the maximum resorptive capacity of the applied cells, such as cells from osteoporotic patients. The model using Neutral medium provides a near-equilibrium situation under control of OB-OC communication, and open to external manipulation. Using Neutral medium, the model could be used to study the effect of drugs or other biochemical compounds on formation and resorption of donor cells, such as cells from osteoporotic patients.

OCs exceeded their expected lifespan. The expected *in vitro* lifespan of OCs is approximately 2 weeks [50]. This roughly correlates with the resorption and TRAP data of the OC medium group, as the TRAP release into the supernatant was decreasing at this point. Remarkably though, the TRAP release was still increasing at d 21 in the Neutral medium group following a linear trend. This suggests that the 2-week lifespan was exceeded in this study. Others have recently shown similar findings. Jacome-Galarza *et al*. showed that parabiont labelled OC were seen up to 24 weeks after parabiont separation [51]. They propose that OC are long-lived but need to acquire new nuclei from circulating blood cells. McDonald *et al*. showed that OC can ‘recycle’ into smaller cells called osteomorphs that can relocate through the bloodstream and re-fuse in the presence of soluble RANKL [52]. Both support the notion that OC can, under some circumstances, indeed die or disappear after 2 weeks, but that with the proper environment it is possible to have active OCs in co-culture for longer than 2 weeks. As in our 2D results, we have shown that in the 3D co-culture monocyte-like cells are present during the entire culture duration. These cells could serve as nuclei-donors as described by Jacome-Galarza to prolong the life of existing OCs and could contain next to monocytes also osteomorphs as described by McDonald to generate new OCs elsewhere in the constructs. These results underline that there is still much to be investigated about the lifespan of OCs.

A higher seeding density of monocytes led to more TRAP release, but not always proportionately. Seeding density has been shown to affect the extent of intercellular communication, maturation and the fusion towards OCs [28]. The TRAP results from the neutral medium groups were as expected; a more than 50 % increase in TRAP release with a peak that occurred earlier in the 1.5 M monocytes group. The increased monocyte seeding density likely also facilitated an increase in cell-fusion events [53] although these were not directly measured in this study. The groups cultured in OC medium did not respond the same way. The increased TRAP release by the 1 M monocytes group was as expected, but the 1.5 M monocytes group did not increase its TRAP release proportionally. Instead, the 1 M and 1.5 M monocytes groups released almost equal amounts of TRAP, and coincidently approximately similar amounts as the 1.5 M monocytes group in Neutral medium. This could indicate that there is a maximum amount of TRAP that these cells can release, and that this amount was reached in both media. It could also mean that there was a maximum to the number of cells close to the surface of the constructs, and that remaining cells did not attach, detached again, or migrated further into the construct where TRAP diffusion into the culture supernatant could be more difficult [54,55].

Seeding density affects net resorption in OC medium, but not in Neutral medium. After an initial differentiation phase from monocytes into OCs, resorption of mineralized volume exceeded formation in the OC medium groups. Interestingly, this switch was observed a little earlier in the 1 M monocytes group than in the 1.5 M group but continued for much longer in the 1.5 M group. In the Neutral medium group, resorption was less apparent, although there was resorption in the 1 M seeded group. One explanation of the higher resorption at lower seeding density group would be that the limited amount of osteoclastogenic signaling molecules released by OB were shared by a higher number of seeded cells of which fewer achieved the necessary threshold to differentiate into resorbing OCs [53], although TRAP results do not directly support this hypothesis. Furthermore, the earlier shown mineralized matrix deposition could be masking low the levels of resorption, essentially creating again an equilibrium situation in tissue remodeling. This indicates that using a higher monocyte seeding density in OC medium leads to a less-than-proportionate increase in TRAP release and possibly an increase in resorption. Using a higher seeding density with OC medium could be useful if a model is needed that favors resorption above formation. Using a higher seeding density with Neutral medium increases TRAP release proportionally without forcing the model into a state of net resorption, which could be useful to study the reaction of OC in a state of equilibrium. However, these conclusions are only valid for cells of this particular donor, because there may be a large variation in activity and resorption between donors [56]. Until models are further qualified and validated to work with variable cell numbers and cells with unknown activity, it is strongly recommended to use well characterized monocytes [57].

The 3D *in vitro* OB-OC co-culture has certain limitations. Because mature human OBs and OCs cannot easily be extracted in meaningful numbers for studies such as these from healthy donors, they must be generated from precursors. *In vitro* differentiation of these precursors is largely dependent on using the correct supplements and media, but these are different for OBs and OCs. Differentiation within the co-culture is thus a compromise between multiple media, and an optimal medium has not been determined yet [58]. Osteoblasts, with their lifespan of up to 200 d, are not a limiting factor for these studies [59]. However, OCs have a life limited life expectancy, effectively limiting the duration of the co-culture. Although others have shown that this life expectancy may be oversimplified [51,52], currently the only practical way to extend the co-culture duration is by introducing ‘fresh’ cells during the co-culture. Increasing resolution (decreasing voxel size) of μCT measurements results in more accurate measurements, at the cost of increased scanning duration and radiation exposure of the cells which may affect both OBs [60] and OCs [61] and their function, especially when performing repeated measurements on the same cells. This study used MSCs and monocytes from two different donors. Ideally, both MSCs and monocytes should come from the same donor [62] to mimic a single person’s response with the model. While MSCs can be expanded [63,64] *in vitro*, monocytes cannot [65]. This limits the number of cells that can be used per blood donation/experiment. While the current study focused primarily on mineralized matrix remodeling and the role of OC therein, OBs were only investigated to a lesser extent and osteocytes were omitted completely. Finally, the model has not been validated by comparison with the *in vivo* situation [66].

Future research should focus on validating [67] and refining the herein presented 3D co-culture model and three media to better reflect specific states of health and disease [68], and make it applicable to accept varying cell yields and cells with varying (osteoclastic) activity. One way to validate the model could be to use cells from a single patient and investigate the predictive capacity of the model by verifying if their cells *in vitro* respond similarly to those *in vivo* [66] when exposed to for example treatment options for osteoporosis such as bisphosphonates. For using such a model as a predictive tool in a clinical setting it would have to be developed into a faster, lower-maintenance and higher-throughput tool, because the time, effort and cost at this point make it unsuitable for routine clinical prediction [69]. Points of attention would be to reduce the construct size, the necessary number of cells and total culture duration, and using automated culture and analysis techniques. All taken together, these improvements could pave the way to develop this 3D OB-OC co-culture model into a valuable tool for fundamental research, drug development and personalized medicine.

## Conclusion

This study shows that the current 3D OB-OC co-culture model can be tuned towards pronouncing either matrix deposition, matrix resorption, or a state of equilibrium by applying one of three culture media. OB medium resulted in continued matrix deposition overshadowing any ongoing resorption, while OC medium forced the differentiation of monocytes towards OCs and resulted in resorption after a period of continuing mineralization. Neutral medium contained neither the osteogenic nor osteoclastogenic supplements and was shown to be closely mimicking a situation of equilibrium, facilitating the study of intricate cell-cell interaction and the result thereof on resorption and formation. The 3D OB-OC co-culture model can be used with either of the three media as an *in vitro* co-culture model of human bone formation and resorption for various applications in fundamental research, drug development and personalized medicine.

**S1_Figure_unenhanced.** Original SEM images that were used for image colorization.

**S2_Dataset_raw_data.** All data as used to create the figures in this publication.

